# Reprogramming of 24nt siRNAs in rice gametes

**DOI:** 10.1101/670463

**Authors:** Chenxin Li, Hengping Xu, Fang-Fang Fu, Scott D. Russell, Venkatesan Sundaresan, Jonathan I. Gent

## Abstract

Gametes constitute a critical stage of the plant life cycle, during which the genome undergoes reprogramming in preparation for embryogenesis. Here we characterized the small RNA transcriptomes of egg cells and sperm cells from rice to elucidate genome-wide distributions of 24nt siRNAs, which are a hallmark of RNA-directed DNA methylation (RdDM) in plants and are typically concentrated at boundaries of heterochromatin. We found that 24nt siRNAs were depleted from heterochromatin boundaries in both gametes, reminiscent of siRNA patterns in DDM1-type nucleosome remodeler mutants. In sperm, 24nt siRNAs were spread across broad heterochromatic regions, while in eggs, 24nt siRNAs were concentrated at a smaller number of heterochromatic loci throughout the genome, which were shared with vegetative tissues and sperm. In both gametes, patterns of CHH methylation, typically a strong indicator of RdDM, were similar to vegetative tissues, although lower in magnitude. These findings indicate that the small RNA transcriptome undergoes large-scale re-programming in both male and female gametes, which is not correlated with recruitment of DNA methyltransferases in gametes and suggestive of unexplored regulatory activities of gamete small RNAs in seeds after fertilization.

## INTRODUCTION

In sexual reproduction in angiosperms, cells transition from vegetative to reproductive fates to produce spore mother cells, which undergo meiosis to produce haploid spores, which then undergo mitosis to produce gametes and other cells of the gametophytes. The development of reproductive cells is marked by multiple sex-specific changes in chromatin structure [reviewed in (Wang and Köhler 2017)]. For example, histone H1 and the centromere-specific histone variant CENH3 are depleted from the megaspore mother cell and the egg (Ingouff et al. 2010; She et al. 2013; Ingouff et al. 2017). Heterochromatin is decondensed in the central cell, which gives rise to the endosperm (Pillot et al. 2010; Yelagandula et al. 2014; Ingouff et al. 2017). A similar phenomenon occurs in the pollen vegetative nucleus (the companion cell to sperm, not to be confused with somatic vegetative tissues) (Schoft et al. 2009; Ingouff et al. 2010; Mérai et al. 2014; Hsieh et al. 2016). Most striking is the compaction of sperm chromatin, likely related to deposition of a set of histone variants including an H3 variant [reviewed in (Borg and Berger 2015)]. Specific activity of some types of euchromatic retrotransposons in the male germline in maize might also reflect male-specific chromatin changes (Dooner et al. 2019).

While DNA methylation patterns in general are transmitted stably across generations, there is also evidence for both loss and gain of methylation in specific sequence contexts and cell types in reproduction [reviewed in (Gehring 2019)]. Of particular interest is RNA-directed DNA methylation (RdDM) because of its potential to be erased and re-established de novo by siRNAs [reviewed in (Cuerda-Gil and Slotkin 2016)]. Methylation in the CHH context (mCHH), where H is A, C, or T, is a strong indicator of RdDM in rice [but not in Arabidopsis (Zemach et al. 2013; Stroud et al. 2014; Niederhuth et al. 2016; Tan et al. 2016)]. Live-cell analysis of transgene-driven expression of fluorescent proteins fused to methyl binding domains in Arabidopsis reveals reduced signal for mCHH in megaspore mother cells, microspores, and in sperm, and for mCG in eggs (Ingouff et al. 2017). Whole-genome bisulfite sequencing (WGBS) has been limited by the difficulty of obtaining sufficient quantities of pure reproductive cell types, but methylomes have been produced from sperm and pollen vegetative cell in both rice and Arabidopsis (Calarco et al. 2012; Ibarra et al. 2012; Hsieh et al. 2016; Kim et al. 2019) and from egg cells in rice (Park et al. 2016). Multiple differences are apparent between cell types by WGBS, including in mCHH. In Arabidopsis, sperm has reduced mCHH relative to pollen vegetative cells (Calarco et al. 2012; Ibarra et al. 2012; Hsieh et al. 2016; Walker et al. 2018). The majority of mCHH in Arabidopsis pollen vegetative nuclei is primarily due to activity of the chromomethyltransferase CMT2 rather than RdDM (Hsieh et al. 2016; Borges et al. 2018). In both Arabidopsis and rice sperm, mCHH and mCHG are partially dependent on demethylation of CGs of corresponding loci in the pollen vegetative nucleus, suggesting that chromatin changes in the pollen vegetative nucleus facilitates expression of siRNAs that are transferred into sperm to direct DNA methylation (Calarco et al. 2012; Ibarra et al. 2012; Kim et al. 2019).

Sequencing small RNAs from Arabidopsis whole pollen and isolated sperm indicates that 24nt siRNAs typical of RNA polymerase IV (pol IV) activity are the most abundant length, but 21nt and 22nt siRNAs are increased in abundance in sperm relative to vegetative tissues (Slotkin et al. 2009; Borges et al. 2018). This increase in the ratio of 21 and 22nt siRNAs to 24nt siRNAs in Arabidopsis pollen is typical of siRNAs in vegetative tissues in mutants with compromised DNA methylation (Creasey et al. 2014; McCue et al. 2015; Corem et al. 2018; Fu et al. 2018; Long et al. 2018; Tan et al. 2018). The Arabidopsis pollen vegetative nucleus, however, does not have reduced DNA methylation characteristic of *ddm1* mutants, nor does sperm except in mCHH (Calarco et al. 2012; Ibarra et al. 2012; Hsieh et al. 2016; Walker et al. 2018).

Mutants with compromised RdDM have been found to affect multiple post-fertilization processes, such as zygotic genome activation (Autran et al. 2011), genome dosage sensitivity in endosperm (Erdmann et al. 2017; Borges et al. 2018; Martinez et al. 2018), imprinting [reviewed in (Anderson and Springer 2018)], and seed development (Grover et al. 2018). There are currently no available small RNA transcriptomes from sperm cells other than Arabidopsis, and none from egg cells for any plant. Due to the small size of the sperm cell relative to the egg cell (∼1/1000 the volume), the transcriptome of the fertilized egg is primarily determined by the egg cell (Autran et al. 2011; Del Toro-De León et al. 2014; Anderson et al. 2017; Zhao et al. 2019). Given their likely importance both pre- and post-fertilization, here we have sequenced small RNAs from both isolated sperm cells and egg cells of rice, compared expression patterns of different classes of small RNAs, and investigated the relationship of 24nt siRNAs to genome-wide DNA methylation in the gametes.

## RESULTS

We prepared and sequenced small RNA Illumina libraries from seedling shoots, isolated sperm cells, isolated egg cells, and whole ovaries (after manual removal of egg cells) of rice variety *Kitaake*. For each cell or tissue type, we prepared at least three biological replicates (**Supplemental Table S1**). We also included several published small RNA libraries from vegetative tissues: wild-type root (Shin et al. 2018), and *ddm1a ddm1b* double mutant leaf (which we refer to simply as *ddm1*) and its wild-type control (Tan et al. 2016; Tan et al. 2018). We aligned all reads of 20 to 25 nucleotides in length to the MSU7 reference genome assembly (Kawahara et al. 2013).

### Distinct miRNA expression patterns in gametes

We first looked at microRNA (miRNA) expression across different sample types and produced a complete list of miRNA abundances (**Supplemental Dataset 1**). miRNAs are predicted to be more cell-autonomous than siRNAs (Grant-Downton et al. 2013), and since mRNA expression is globally different between egg, sperm, and seedling (Russell et al. 2012; Anderson et al. 2013; Anderson et al. 2017), we also expected regulators of gene expression, such as miRNAs, to be differentially expressed across sample types. We first performed principal component analysis (PCA, **Fig. 1A**) to examine global differences of miRNA expression across samples. On a PCA plot, the sample types were clearly separated. Vegetative vs. floral/reproductive samples were separated along the first PC axis, which explained ∼37% of variance in our miRNA expression dataset. Male and female samples were separated along the second PC axis, which explained ∼12% of the variance in our miRNA expression dataset. To discover which individual miRNA drove the variation between sample types, we performed hierarchical clustering and detected multiple miRNA clusters that distinguished seedling shoot from both gametes, as well as clusters that were enriched in one gamete but not the other (**Supplemental Fig. S1A** and **Supplemental Table S3**). We also found that most clusters correlated with either one of the PCs (**Supplemental Fig. S1B**). Thus, the separation of samples on PC plots are likely driven by miRNA clusters that have distinct expression patterns across samples. miR159 family members were the most abundant miRNAs, especially in egg and ovary (**Fig. 1B**). In fact, five copies of miR159 accounted for 76% of the total miRNAs in egg and 90% in ovary. Seedling shoot had an intermediate level, 46%, and sperm had 8%. While two family members, miR159a.1 and miR159b, share an identical mature miRNA sequence that cannot be distinguished, the rest of the family members had distinct expression patterns (**Fig. 1B**). miR159a.2 was unique to root, while miR159c, d, and e were enriched in egg, and miR159f was enriched in seedling shoot and egg. In Arabidopsis, miR159 family members repress MYB transcription factors and are expressed in multiple tissues including sperm, and paternal miR159 regulates nuclear division in endosperm (Allen et al. 2010; Zhao et al. 2018).

**Figure 1.**
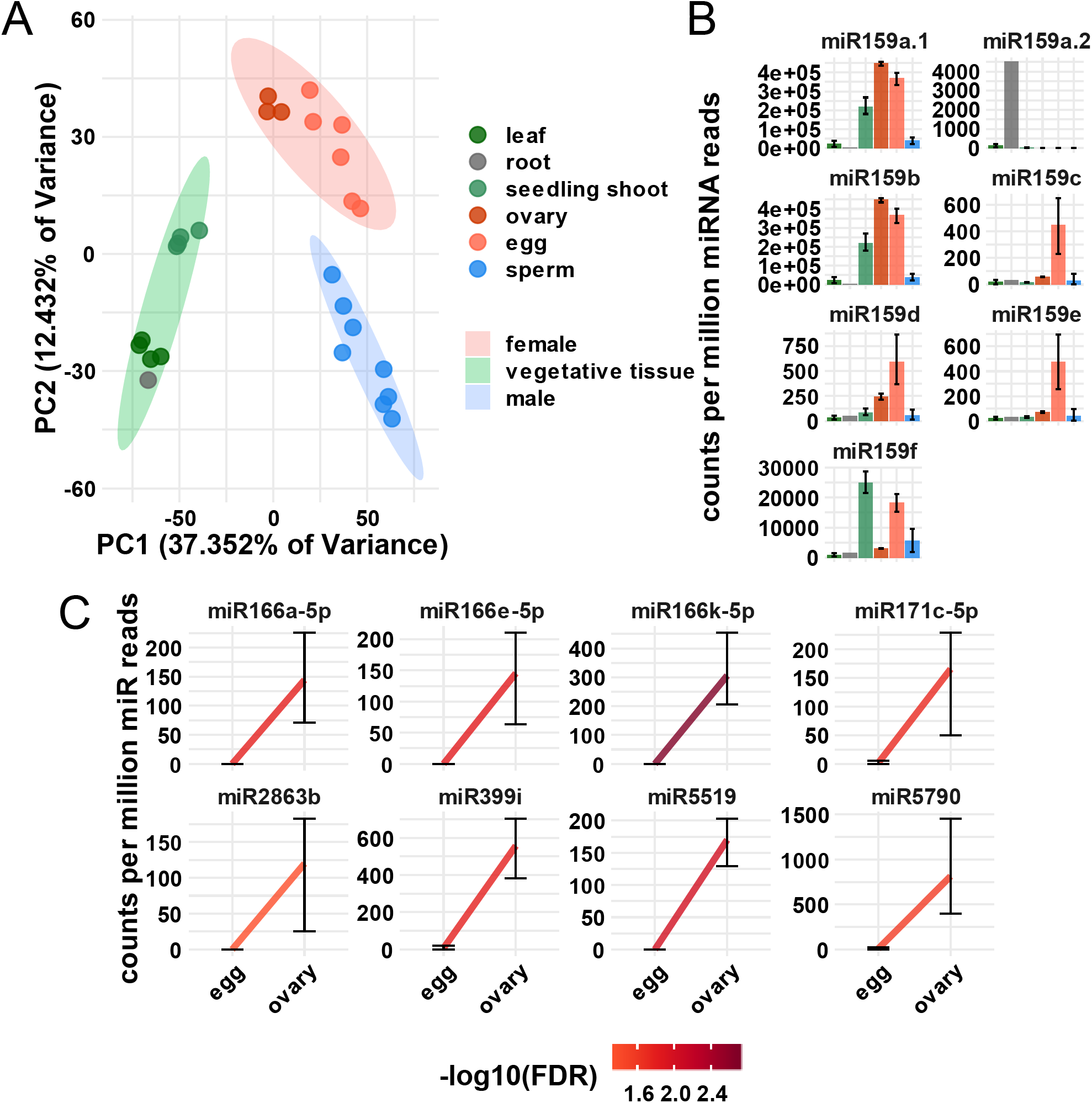
miRNA expression patterns. **A:** Principle Component (PC) plot of sample types by miRNA expression pattern. **B:** miRNA159 family expression pattern. Y-values are relative to the total number of miRNA reads in each sample. Error bars are 95% confidence intervals for samples with two or more biological replicates. Color code is the same as in **A. C:** Representative differentially expressed miRNAs between egg and ovary. Differential expression was determined by 2-fold change and FDR (false discovery rate) < 0.05 cutoffs. Y-values are relative to the total number of miRNA reads in each sample. Error bars are 95% confidence intervals. Color of line reflects −log10 of FDR.

The manual dissection of ovaries to isolate egg cells raises the concern of releasing small RNA from ovary into solution and being carried along with egg cell small RNAs into sequencing libraries. Three lines of evidence indicated RNA contamination from ovary in egg cells was minimal. First, our egg or zygote isolation method has previously been shown to have little mRNA contamination from maternal tissue (Anderson et al. 2017; Li et al. 2019). Second, we performed differential expression analysis of the miRNAs, and found that eight miRNAs had high levels in all three replicates of ovary but were nearly undetectable in all six replicates of egg cells (**Fig. 1C**). Third, we did mock egg cell isolations in which we dissected ovaries using the same protocol, but we only collected cell-free solution rather than egg cells. By qPCR quantification, library preparation from the mock samples yielded only about 5% of the number of library molecules as an actual egg cell small RNA library after 20 cycles of PCR amplification (**Supplemental Fig. S2**). Thus, major contamination from ovary tissue to isolated egg cell samples was very unlikely. Similarly, our sperm isolation method has previously been shown to have little mRNA contamination from surrounding floral tissues or pollen (Li et al. 2019).

### Distinct small RNA composition across gametes and vegetative tissues

We noticed that small RNA compositions in gametes were distinct from general vegetative tissues, such as seedling shoot, most noticeably by the reduction of miRNAs relative to total small RNAs (**Fig. 2A**). In seedling shoot, 10% of the small RNA reads mapped to miRNA loci, as compared to 2.5% in egg and 0.5% in sperm. In addition to miRNAs, the relative abundance of phased secondary siRNAs (phasiRNAs), tRNAs, NOR RNAs, and 5S rRNAs varied across gametes and vegetative tissues. The relative abundance of 21nt and 24nt phasiRNAs clearly differentiated egg and sperm. While both classes were rare in egg cells relative to seedlings, 21nt phasiRNAs were even rarer in sperm (**Fig. 2A**). In contrast, 24nt phasiRNAs were greater than 30-fold more abundant in sperm than other sample types. In maize, pre-meiotic anthers accumulate 21nt phasiRNAs, whereas meiotic anthers accumulate 24nt phasiRNAs that persist into pollen (Zhai et al. 2015b). Similar processes may occur in rice, which may explain the abundance of 24nt phasiRNAs in rice sperm, since 24nt phasiRNAs have been previously demonstrated in rice meiotic inflorescences (Johnson et al. 2009).

**Figure 2.**
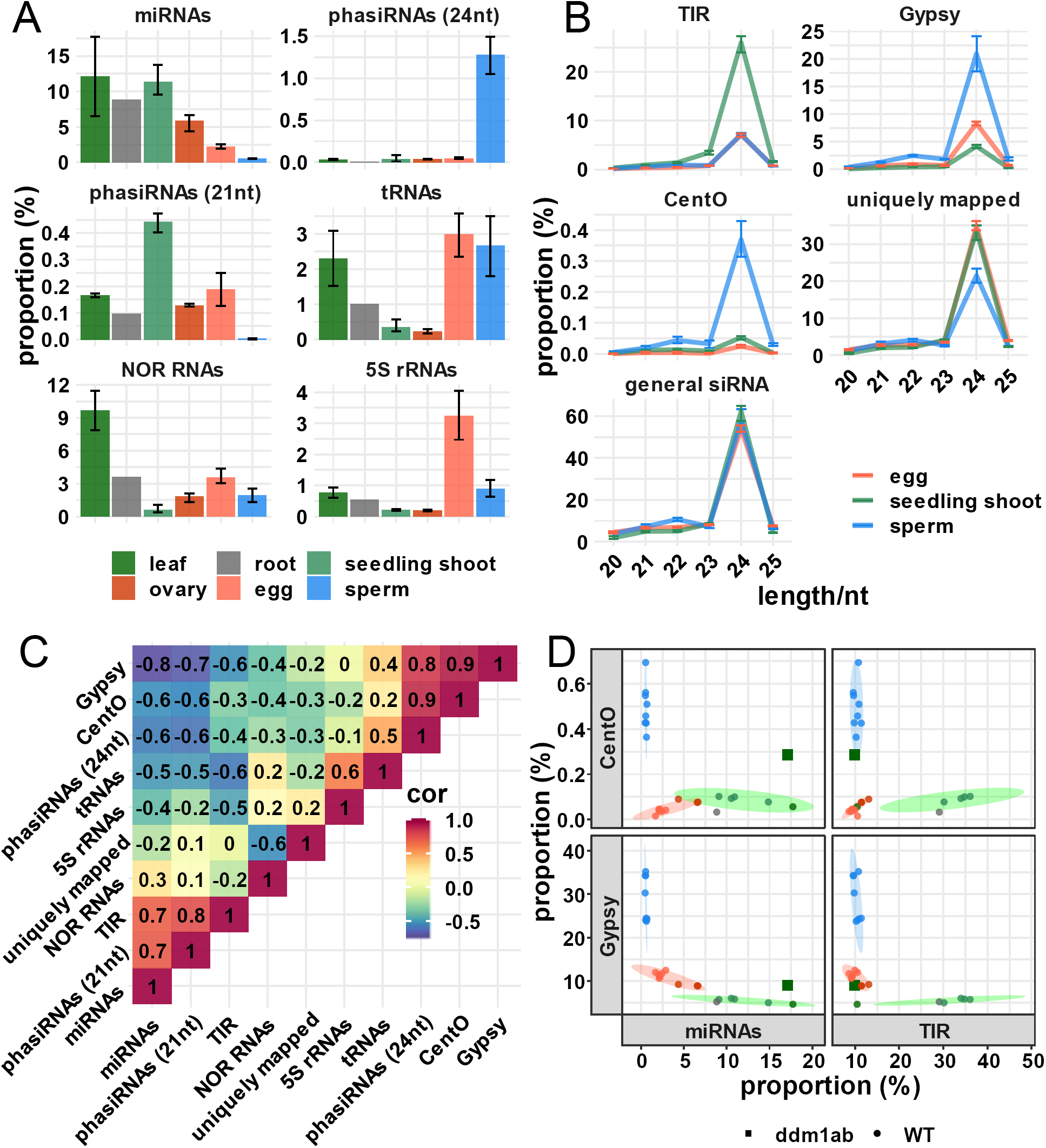
Distinct small RNA composition in gametes. **A:** small RNA composition across sample types. Y-axis values are relative to the total number of reads that mapped to the genome. For samples with two or more biological replicates, error bars are 95% confidence intervals. **B:** siRNA abundance by length and category. “siRNAs” refers to all 20-25nt small RNAs that mapped to the genome but were not included in any category in **A**. “Uniquely-mapping” are the subset that map with a MAPQ value of at least 20. “*Gypsy*” are the subset that overlap with an annotated *Gypsy* element, and “TIR” are the subset that overlap with an annotated DNA transposon of the *Tc1/Mariner, PIF/Harbinger, Mutator* or *hAT* superfamilies. “*CentO*” are the subset that overlap a *CentO* tandem repeat. Y-axis values are the number of siRNAs for each length normalized by the total number of siRNAs. Error bars are 95% confidence intervals. **C:** Correlation heatmap of different small RNA types. Color gradient and text reflect correlation coefficients between a pair of small RNA types across wild-type libraries. **D:** Scatter plots showing sample types clustered based on proportions of specific types of siRNAs or miRNAs. First row: Y-axis is proportion of *CentO* siRNAs. Second row: Y-axis is proportion of *Gypsy* siRNAs. First column: X-axis is proportion of miRNAs. Second column: X-axis is proportion of TIR siRNAs. Color code is the same as **A**.

The most abundant small RNAs in angiosperm vegetative tissues are 24nt long siRNAs that are involved in RNA-directed DNA methylation (RdDM), and to a lesser extent 21 and 22nt siRNAs [reviewed in (Cuerda-Gil and Slotkin 2016)]. We grouped all 20-25nt small RNAs that did not overlap at least 90% of their length with miRNA, phasiRNA, tRNA, NOR RNA, or 5S rRNA loci into a general siRNA category for further analysis. The length distribution of these siRNAs was similar between cell and tissue types, with a strong preference for 24nt (**Fig. 2B**).Parsing out uniquely aligning siRNAs from the total, however, revealed a depletion in uniquely-mapping 24nt siRNAs in sperm relative to seedling shoot and egg cell (**Fig. 2B**). Further parsing out siRNAs that overlapped by at least 90% of their length with retrotransposons of the *Gypsy* superfamily and the centromeric tandemly repeat *CentO* revealed strong siRNA enrichments for these elements in sperm. *CentO* tandem repeats are enriched in centromeres (Nagaki et al. 2003), and *Gypsy* retrotranposons are generally enriched in heterochromatin regions of the genome (Kawahara et al. 2013). siRNAs overlapping with Terminal Inverted Repeat (TIR) DNA transposons of the *PIF/Harbinger, Tc1/Mariner, Mutator* or *hAT* superfamilies were enriched in seedling shoot but were depleted in both sperm and egg. For brevity we refer to siRNAs from these four superfamilies of DNA transposons as simply TIR siRNAs. These transposons are enriched near genes (Han et al. 2013).

Pairwise correlations between different small RNA classes across all wild-type samples revealed that relative abundance of miRNAs, TIR siRNAs, and 21nt phasiRNAs were positively correlated and were high in vegetative tissues, while relative abundance of *Gypsy* siRNAs, *CentO* siRNAs, and 24nt phasiRNAs were positively correlated and were high in sperm. Egg and ovary small RNAs were intermediate between those of vegetative tissues and sperm. The two co-abundant groups of small RNAs were mutually anti-correlated (**Fig. 2C**). Plotting samples by their relative abundances of different small RNA classes showed the same trends and demonstrated that samples clustered according to their tissue type (**Fig. 2D**). We further included in this analysis published small RNA data from *ddm1* mutant leaf tissue (Tan et al. 2016; Tan et al. 2018). *ddm1* leaf diverged from wild-type leaf by a large increase in relative abundance of *CentO* siRNAs, which is more similar to sperm. In addition, *ddm1* leaf also had a higher relative abundance of *Gypsy* siRNAs, which is more similar to both sperm and egg. Thus, differences in small RNA composition, especially differences in siRNAs mapping to repetitive elements, may in part be explained by chromatin alterations in gametes similar to those that occur with loss of DDM1 activity. In summary, these results indicate that 1) large-scale reprogramming of small RNA transcriptomes occurs in reproductive cells, leading to distinct small RNA distributions in male and female gametes, and 2) the changes in siRNA distributions are consistent with differential chromatin structure.

### Distinct genome-wide patterns of 24nt siRNAs in gametes

Consistent with the high abundance of *CentO* and *Gypsy* siRNAs, a genome-wide view of 24nt siRNA abundance revealed that 24nt siRNAs in sperm cells had a striking complementary pattern to that of seedling shoot. Sperm cell 24nt siRNAs were enriched in gene-poor, heterochromatic regions, whereas seedling shoot 24nt siRNAs had the expected pattern for canonical RdDM in vegetative tissues, with enrichment in euchromatic regions that are gene-rich (**Fig. 3A and Supplemental Fig. S3A**). Egg cell 24nt siRNAs showed a pattern distinct from both sperm and shoot, with depletion across most of the genome except at a smaller number loci with no clear relationship to gene density. The distribution of 21 and 22nt siRNAs also did not have a clear relationship to gene density, in neither the gametes nor seedling shoot (**Supplemental Fig. S3A**). Interestingly, 24nt siRNAs from ovary showed a nearly identical pattern with egg, but the 21 and 22nt siRNAs pattern diverged between ovary and egg (**Supplemental Fig. S3A**), as did some miRNAs (**Fig. 1A**). Consistent with the genome-wide analysis, metagene plots of the distribution of 24nt siRNA relative to genes revealed a depletion of 24nt siRNAs both upstream and downstream of genes in both egg and sperm (**Fig. 3B**), corresponding to where TIR transposons are enriched in the genome, with the exception of the CACTA superfamily (Han et al. 2013).

**Figure 3.**
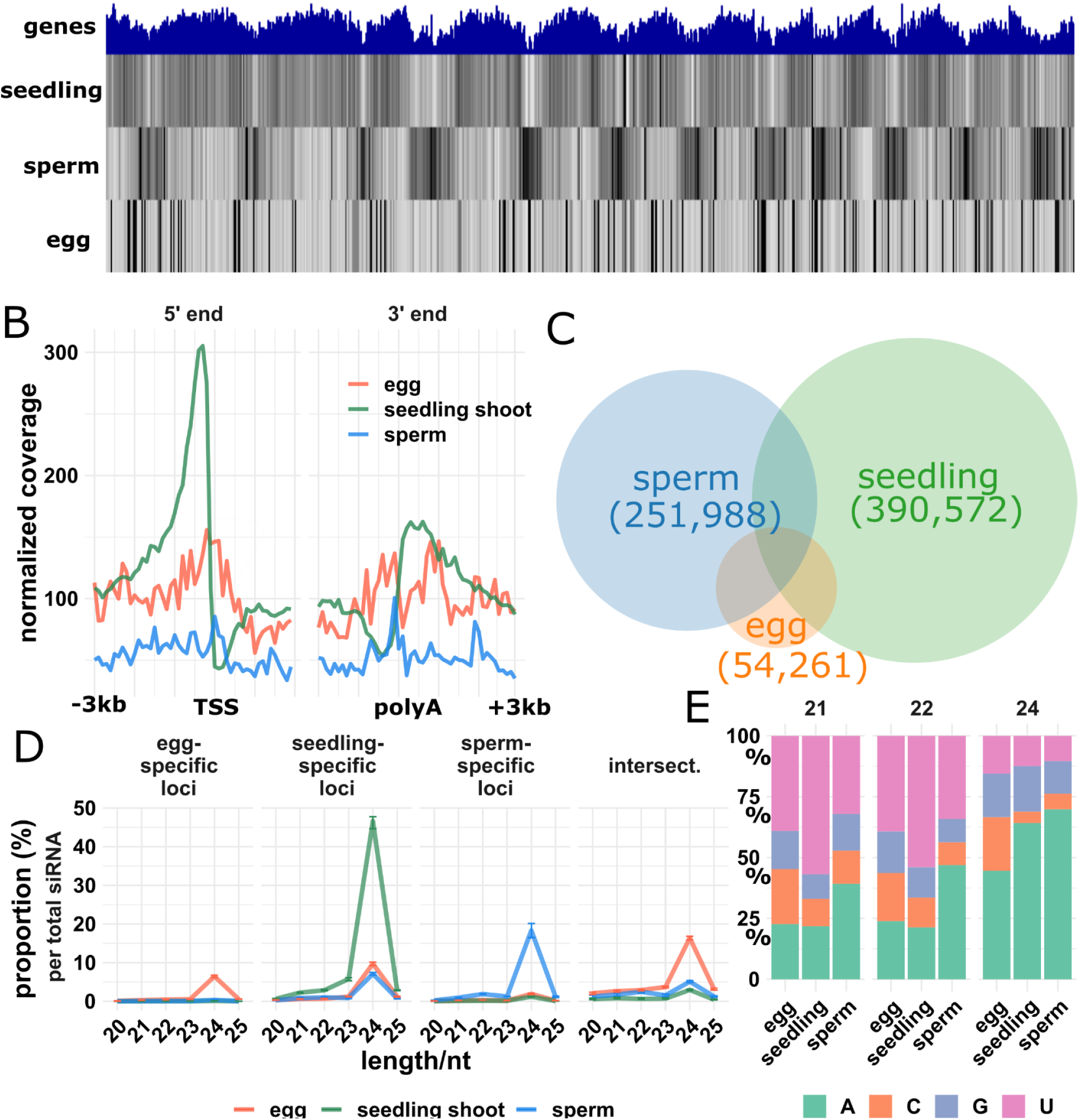
Distinct genome-wide small RNA distribution in gametes. **A:** genome-wide view of 24nt siRNAs. Top track: gene density; second to fourth tracks: seedling shoot, sperm and egg, respectively; Chr: chromosome. **B:** Metagene plot for 24nt siRNAs. Plots indicate 24nt siRNA coverage with 100bp resolution from 3KB upstream to 3KB downstream of genes, normalized per 1000 total siRNAs. Major and minor tick marks indicate 1000bp and 500bp intervals, respectively. TSS: Transcription start site; poly A: polyadenylation site. **C:** Venn diagram showing number and overlap of 24nt siRNA loci. The genome was divided into 100bp intervals and categorized as 24nt siRNA loci based on coverage of 24nt siRNAs from seedling shoot, sperm, or egg. **D:** siRNA abundance at 24nt siRNA loci. Sample-specific loci were only categorized as such in one tissue. “Intersect.” refers intersection loci, those categorized as 24nt siRNA in all three tissues. Y-values are the number of siRNAs for each length divided by the total number of siRNAs that mapped to the genome. Error bars are 95% confidence intervals. **E:** 5’-nucleotide preferences in 24nt siRNAs. The stacked bar charts show the percent of 24nt siRNAs from the sample-specific 24nt siRNA loci categories that begin with each nucleotide.

To identify specific loci associated with siRNAs in each tissue, we divided the genome into non-overlapping 100bp loci and counted the number of overlapping siRNAs per 50 million total siRNAs from each sample. Any locus that had at least 25 overlapping 24nt siRNAs that spanned at least 50bp of DNA we categorized as a 24nt siRNA locus. In seedling shoot, 390,572 loci met these criteria, in sperm 251,988, and in egg 54,261 (**Fig. 3C**). Consistent with the complementary patterns of 24nt siRNA loci in sperm and seedling shoot in the whole genome view of siRNA abundance (**Fig. 3A**), only 15% of the sperm loci were shared with seedling shoot loci. In contrast, 45% of the egg loci were shared with seedling shoot. To determine how well each set of 24nt siRNA loci represented the siRNAs from each tissue, we counted the number of 24nt siRNAs that overlapped with them (**Table 1**). In seedling shoot, 80% of 24nt siRNAs overlapped with seedling shoot 24nt siRNA loci. The other 20% of 24nt siRNAs mapped to locations with insufficient numbers of siRNAs to meet our threshold values to qualify as 24nt siRNA loci. Only 50% of sperm 24nt siRNAs overlapped with sperm 24nt siRNA loci, indicating that 24nt siRNAs were less concentrated at specific loci in sperm than in seedling shoot. Egg cells showed an intermediate makeup, with 63% of 24nt siRNAs overlapping with egg 24nt siRNA loci. The loci that were shared between all three tissues we called intersection 24nt siRNA loci, while the loci that were unique to one tissue we called either seedling-specific, sperm-specific, or egg-specific. Taking into account the number of loci in each category relative to the size of the genome revealed that intersection loci were more strongly enriched for 24nt siRNAs than sample specific loci in all three tissues (**Table 1**). 24nt siRNAs were the dominant siRNAs in tissue-specific and intersection loci, but other siRNA lengths were also abundant, particularly in intersection loci (**Fig. 3D, Supplemental Fig. S4**). Counting overlaps between 24nt siRNA loci and repetitive elements and measuring distances between 24nt siRNA loci and nearest genes revealed expected trends of seedling-specific 24nt loci being near genes and overlapping with TIR transposons, typical of canonical RdDM. (**Supplemental Fig. S5A**).Gamete-specific loci were farther from genes and overlapped more with *Gypsy* retrotransposons, whereas intersection loci were intermediate between gamete-specific and seedling-shoot, but more similar to seedling-specific in their distances to nearest genes (**Supplemental Fig. S5**).Taken together, these data reveal that sperm and egg cells have distributions of siRNAs that are uncharacteristic of canonical RdDM in rice and that are distinct from each other. The sperm cell is complementary to vegetative tissues in its enrichment of siRNAs in large numbers of heterochromatic loci, while the egg cell has a different pattern entirely, with large number of siRNAs in a small number of loci that tend to be shared in sperm and vegetative tissues.

**Table 1:** Number of 24nt siRNA loci and proportion of siRNAs that overlapped them in seedling shoot, sperm, and egg. Proportion of 24 siRNAs is the number of 24nt siRNAs that overlapped with the loci divided by the total number of 24nt siRNAs in the sample. Enrichment of 24nt siRNAs in loci is the proportion 24nt siRNAs that mapped to loci normalized by the number of loci in the genome.

| 24nt siRNA loci type | Number of loci | Proportion of 24nt siRNAs in loci |  |  | Enrichment of 24nt siRNAs in loci |  |  |
| --- | --- | --- | --- | --- | --- | --- | --- |
|  |  | seedling | sperm | egg | seedling | sperm | egg |
| seedling | 390572 | 0.80 | 0.28 | 0.50 | 7.7 | 2.7 | 4.8 |
| sperm | 251988 | 0.11 | 0.50 | 0.49 | 1.6 | 7.5 | 7.3 |
| egg | 54261 | 0.11 | 0.14 | 0.63 | 7.5 | 9.9 | 43.0 |
| seedling-specific | 338826 | 0.65 | 0.12 | 0.16 | 7.2 | 1.3 | 1.8 |
| sperm-specific | 197300 | 0.02 | 0.30 | 0.03 | 0.3 | 5.6 | 0.5 |
| egg-specific | 13796 | 0.00 | 0.01 | 0.11 | 0.4 | 1.5 | 28.9 |
| intersection | 11103 | 0.04 | 0.08 | 0.26 | 13.9 | 28.0 | 88.8 |
| sperm-seedling intersection | 38536 | 0.09 | 0.16 | 0.28 | 8.5 | 15.4 | 26.8 |
| sperm-egg intersection | 27255 | 0.04 | 0.13 | 0.45 | 5.9 | 18.0 | 62.1 |
| egg-seedling intersection | 24313 | 0.11 | 0.09 | 0.33 | 16.2 | 13.9 | 50.7 |

To test whether the pattern of 24nt siRNAs in either sperm or egg could be explained by loss of 24nt siRNAs from canonical RdDM loci, we removed all seedling shoot 24nt siRNAs that overlapped with seedling-specific 24nt siRNA loci, then examined the distribution of the remaining seedling 24nt siRNAs in the genome. While this did result in fewer siRNAs near genes, it did not produce the enrichment in heterochromatic regions observed in sperm (**Supplemental Fig. S3B**). Canonical RdDM is indicated by mCHH in addition to 24nt siRNAs, so we also identified putative canonical RdDM loci independently, using mCHH levels in vegetative tissues from a previous study (Tan et al. 2016). We divided the genome into non-overlapping 100bp loci, then identified loci with an average mCHH of at least 5%. 315,368 loci, 8% of the genome, met these criteria, and 53% of seedling shoot 24nt siRNAs overlapped with these loci. After removing seedling 24nt siRNA from mCHH loci, the residual 24nt siRNA in seedling shoot were relatively depleted near genes, but again there was no specific enrichment in heterochromatic regions, as was the case with sperm, nor was there a pattern resembling that of egg cells (**Supplemental Fig. S3B**).

Expression of sperm genes has an atypical distribution, with a smaller number of genes accounting for a larger proportion of mRNA in sperm than egg (Anderson et al. 2013) (**Supplemental Fig. S6A**). Since abundance of 24nt siRNAs and mCHH near genes has been positively correlated with gene expression (Li et al. 2015), we wondered whether highly expressed sperm genes might be enriched for flanking 24nt siRNAs. This was not the case, however. Genes with transcripts per million (TPM) values of greater than 10, which corresponds to the top ∼25% of mRNA expression level in sperm, exhibited similar lack of flanking siRNAs as the total set of genes (**Supplemental Fig. S6B**). We also asked whether differences in siRNAs could be explained by differences in mRNA levels of RdDM factors or DDM1 in egg and sperm. Analysis of published gamete transcriptomes did not reveal any obvious explanation for egg or sperm siRNA patterns, though they did show dramatic differences in mRNA levels for some RdDM factors such as RDR2 (Anderson et al. 2013; Anderson et al. 2017) (**Supplemental Fig. 7**). At least one DDM1 homolog was strongly expressed in both gametes, but in pollen vegetative cell a single homolog was only moderately expressed. Analysis of 5’ nucleotide preferences of gametic and vegetative 24nt siRNAs revealed a strong bias toward a 5’ A and against a 5’ U in seedling shoot and in sperm (**Fig. 3E**), consistent with known pol IV activity (Blevins et al. 2015; Zhai et al. 2015a). The 5’ nucleotide biases were also in egg, but weaker, suggesting that egg cell 24nt siRNAs might also arise from different RNA polymerase activities or processing of siRNA precursors (both pol II and pol IV and multiple dicers and other factors can interact to produce diverse siRNAs (Cuerda-Gil and Slotkin 2016)). In sperm, 21-22nt siRNAs also exhibited a preference for the 5’ A nucleotide, though weaker than that of 24nt siRNAs, consistent with reports that pol IV can produce 21nt siRNAs in Arabidopsis pollen (Borges et al. 2018; Martinez et al. 2018).

Transposon-related mRNAs made up a higher proportion of the total mRNA in sperm than in egg, zygote, or seedling (Anderson et al. 2013; Anderson et al. 2017) (**Supplemental Fig. S8**). This was true both for heterochromatic transposons (represented by LTR retrotransposons of the *Gypsy* superfamily) and euchromatic ones (represented by TIR transposons). Analysis of siRNA and mRNA expression of individual transposon copies is complicated by the limitation in mapping short reads to them uniquely. However, a strong trend was evident in that TE copies with high mRNA expression in gametes tended to have fewer 24nt siRNAs, and vice versa (**Supplemental Fig. S9**). Both *Gypsy* and *TIR* transposons exhibited this relationship, suggesting that 24nt siRNAs might be generally antagonistic to transposon mRNA accumulation.

### Gametes have similar DNA methylation as vegetative tissues

24nt siRNA are key components of RdDM in vegetative tissues. To test whether the unusual genomic distributions of siRNAs in sperm and egg were associated with RdDM, we prepared and sequenced whole genome bisulfite sequencing (WGBS) libraries using a post-bisulfite adapter tagging (PBAT) method from isolated egg and sperm (**Supplemental Table S2**). We also analyzed published PBAT libraries from egg, sperm, central cell, and pollen vegetative cell (Park et al. 2016; Kim et al. 2019). The DNA methylation in all of these cell types resembled typical vegetative tissues in that mCHH was highest near genes (**Fig. 4A, Supplemental Figs. S10 and S11**). Despite the fact that 24nt siRNA distributions were drastically different between egg, vegetative tissues, and sperm, genome-wide distributions of DNA methylation were similar between them, though the magnitude of methylation was variable between cell types and even between experiments. We also prepared libraries from whole ovary, bract (palea and lemma), embryo, and endosperm using a conventional WGBS method **(Supplemental Table S2**). All wild-type conventional libraries revealed the expected pattern of high mCHH flanking genes except mature endosperm (**Fig. 4B**). As with egg, the ovary retained robust mCHH at canonical RdDM loci, despite loss of 24nt siRNAs (**Supplemental Fig. S3A**). We also generated a knockout mutant for *drm2* using CRIPSPR-Cas9 gene editing and prepared libraries from mutant endosperm and embryo. *Drm2* encodes a DNA methyltransferase of the Domains Rearranged Methyltransferase family, which function in RdDM. Near-gene mCHH was almost abolished in *drm2* embryo and endosperm (**Fig. 4B** and **Fig. 5B**), demonstrating that the mCHH peaks near genes require functional DRM2, and consistent with prior work on methylation in *drm2* mutant leaf (Tan et al. 2016).

**Figure 4.**
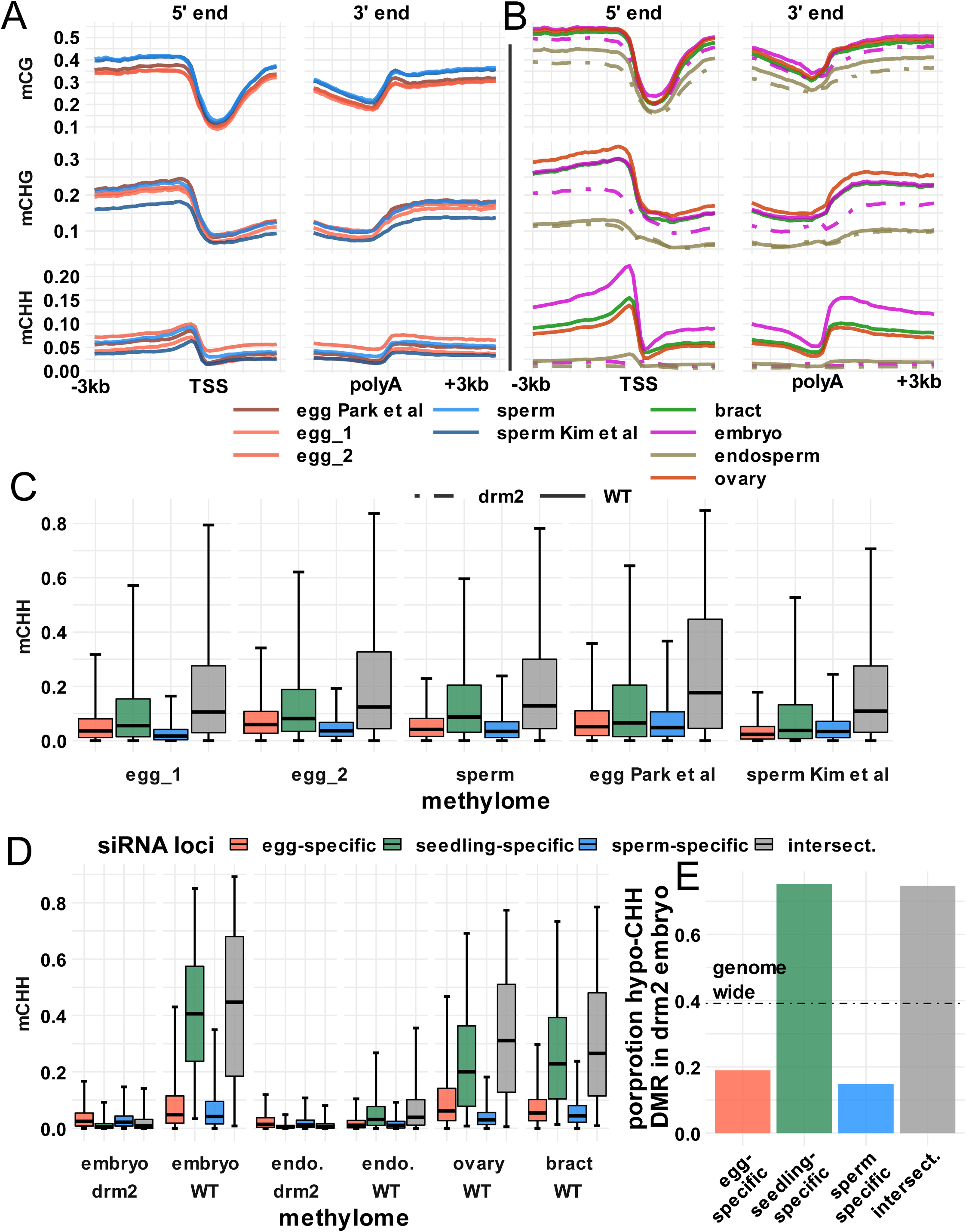
Gametes have similar methylation patterns. **A:** DNA methylation metagene plots for post-bisulfite adapter tagging (PBAT) libraries. Plots indicate average DNA methylation values over 100bp intervals from 3KB upstream to 3KB downstream of genes. Major and minor tick marks indicate 1000bp and 500bp intervals, respectively. DNA methylation is measured as the percent methylated cytosines relative to total cytosines in each sequence context. TSS: Transcription start site; poly A: polyadenylation site. **B:** DNA methylation metagene plot for conventional WGBS libraries, X and Y-axes as in **A. C:** DNA methylation of 24nt siRNA loci for PBAT libraries. Center line is median methylation; box represents interquartile range; whiskers extend to 97.5^th^ percentile and 2.5^th^ percentile, respectively. **D:** DNA methylation of 24nt siRNA loci for conventional WGBS libraries, as in **C**. Color code the same as **C. E:** Proportion of hypomethylated CHH differential methylated regions (DMR) in mature *drm2* embryo relative to wild-type. Dotted line represent genome wide proportion of hypo-CHH DMR. Bract: lemma and palea of rice florets. Endo: endosperm.

**Figure 5.**
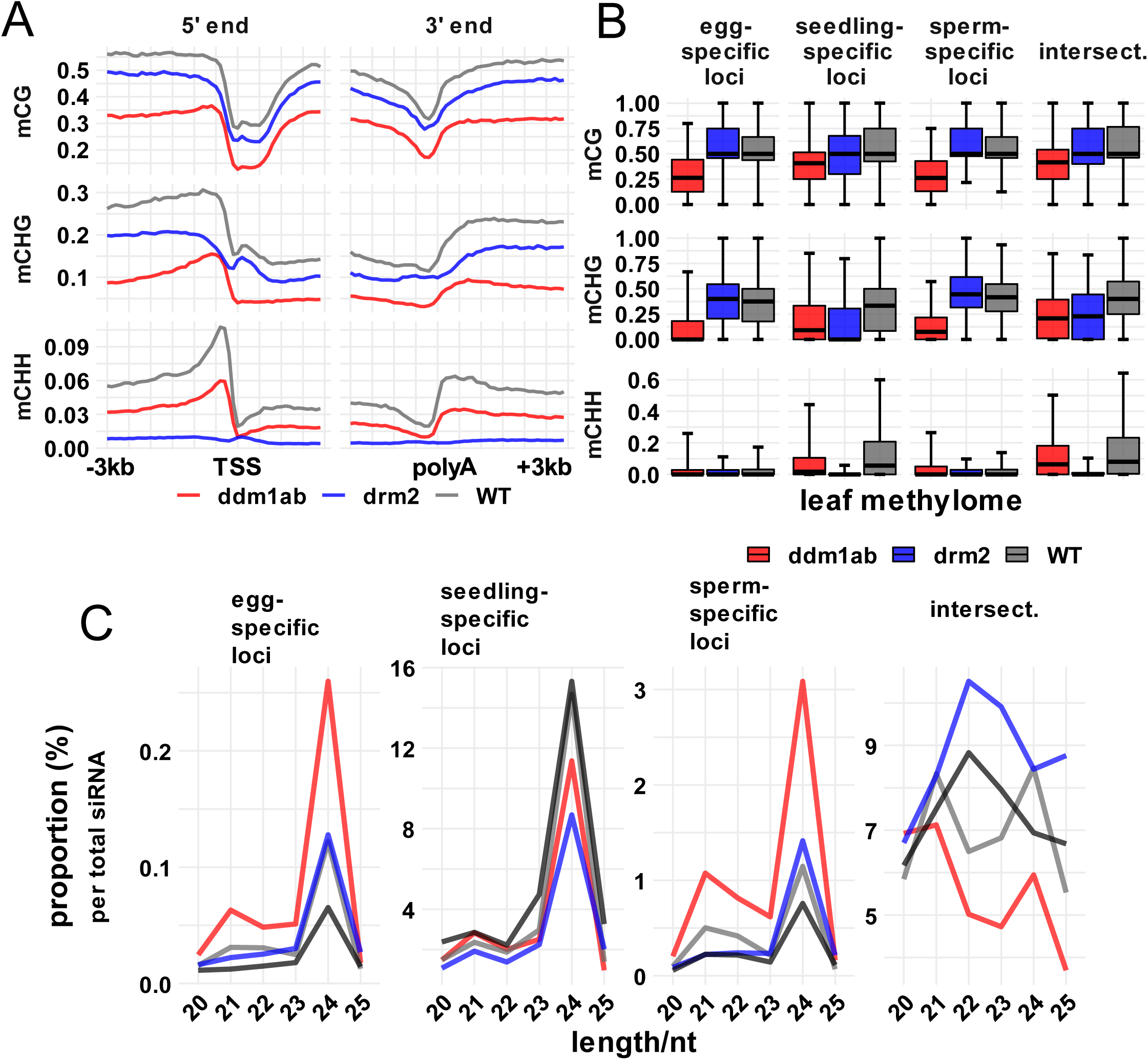
siRNA and methylation profiles in *ddm1* and *drm2* mutant leaf. **A:** DNA methylation metagene plots. Plots indicate average DNA methylation values over 100bp intervals from 3KB upstream to 3KB downstream of genes. Major and minor tick marks indicate 1000bp and 500bp intervals, respectively. DNA methylation is measured as the percent methylated cytosines relative to total cytosines in each sequence context. TSS: Transcription start site; poly A: polyadenylation site. **B:** DNA methylation of 24nt siRNA loci in *ddm1a ddm1b double mutant, drm2* and wild-type leaf methylomes (data from Tan et al 2016). Center line is median methylation; box represents interquartile range; whiskers extend to 97.5^th^ percentile and 2.5^th^ percentile, respectively. **C:** siRNA abundance at 24nt siRNA loci. Y-values are the number of siRNAs of each length divided by the total siRNAs that mapped to the genome.

To test whether 24nt siRNAs directed DNA methylation in gametes, we examined gamete DNA methylation in the tissue-specific 24nt siRNA loci that we identified (see above). All tissue-specific classes of 24nt siRNA loci had high mCG and mCHG in all wild-type samples, including gametes (**Supplemental Fig. S12**). Endosperm was an exception, with reduced mCHG. Only seedling-specific and intersection loci had high mCHH levels though, independently of 24nt siRNAs. In other words, despite the abundance of 24nt siRNAs in egg-specific 24nt siRNA loci, these loci had lower mCHH in egg than seedling-shoot specific loci in egg. The same was true for sperm: sperm-specific 24nt siRNA loci had lower mCHH in sperm than seedling-shoot specific loci in sperm (**Fig. 4C, Fig. 3D**). Intersection loci had the highest mCHH across the four locus categories. This pattern was true not only for egg and sperm, but also for pollen vegetative cell, central cell, and vegetative tissues (**Supplemental Fig. S12A**). Although variation between tissues and even between experiments exists (**Fig. 4A)**, importantly, within-library patterns are reproducible and confirm that redistribution of 24nt siRNAs is not coupled to redistribution of DNA methylation in gametes (**Fig. 4C)**. In all cases, differences between gamete-specific loci and seedling-specific loci were highly significant (p < 10^-5^, Tukey test using generalized linear model with logit link). In addition, *drm2* mutations abolished mCHH in seedling-specific 24nt loci and intersection 24nt loci in mature embryo and leaf, further confirming that these are canonical RdDM loci (**Fig. 4D, Fig 5B**). We also confirmed our inferences by counting the proportion of differentially methylated regions (DMRs) in the CHH context between mature WT and *drm2* embryo across 24nt siRNA loci categories (**Fig. 4E**). Genome-wide, around 40% of the eligible loci (loci that had sufficient coverage to be included in DMR analysis) were hypomethylated for mCHH in *drm2* embryo. Seedling-specific and intersection loci were enriched for hypo-mCHH DMR (∼80%), while egg and sperm-specific loci were depleted for hypo-mCHH DMR (∼ 20%). Hypergeometric tests detected significant enrichment or depletion for DMR in these sets of loci, respectively (p value = 0 for all four categories).

### Gamete siRNAs marked genomic regions normally methylated by DDM1 but not DRM2

The loss of siRNAs from TIR DNA transposons and gain from LTR retrotransposons in gametes is reminiscent of *ddm1* mutant phenotypes in vegetative tissues in diverse angiosperms (Creasey et al. 2014; McCue et al. 2015; Corem et al. 2018; Fu et al. 2018; Long et al. 2018; Tan et al. 2018). In *ddm1* mutants and other mutants that reduce heterochromatic DNA methylation, RNA polymerases may gain access to DNA to which they would not normally have access. The leaf phenotype of a *ddm1* double mutant in rice includes substantial loss of mCHH and siRNAs flanking genes, coupled with increased DRM2 activity and siRNAs in heterochromatic regions (Tan et al. 2016; Tan et al. 2018). The reported differences in siRNAs between wild-type and *ddm1* mutant leaf were not as severe as the differences we found between wild-type seedling shoot and wild-type gametes. Since the magnitude of such differences can be influenced by analysis methods, we reexamined the published *ddm1* small RNA and methylation data in parallel with our gamete data. We also included published *drm2* mutant data from the same study as the *ddm1* mutant methylation data (Tan et al. 2016). Our analysis of methylation and siRNAs confirmed the prior study in that 24nt siRNAs were reduced in gene flanks in the *ddm1* mutant, but not as severely as in gametes (**Supplemental Fig. S13**). Unlike gametes but in agreement with the previous study, the *ddm1* mutant also had a strong reduction in methylation (∼50% lower), including mCHH, in gene flanks in leaf (**Fig. 5A**). The *drm2* mutant had stronger reduction in mCHH but weaker reduction in mCG and mCHG near genes, as expected for a factor directly involved in RdDM (**Fig. 5A**).

To investigate whether gamete-specific 24nt siRNA loci might be targets of DDM1 activity, we also evaluated DNA methylation in the mutants for each set of 24nt siRNA loci. We found severe reductions in methylation—particularly in mCHG—in *ddm1* mutants (p < 10^-5^, Tukey test using generalized linear model with logit link), but not in *drm2* mutants, at sperm-specific and egg-specific 24nt siRNA loci (**Fig. 5B**). At these loci, the spread of mCHH values was more variable in the *ddm1* mutant than in wild-type and thus difficult to confidently determine the overall trend. The seedling shoot-specific and intersection 24nt siRNA loci were strongly dependent on DRM2 for mCHH methylation (p < 10^-5^, Tukey test using generalized linear model with logit link), consistent with canonical RdDM activity. The relative abundance of 24nt siRNAs increased about twofold in gamete-specific 24nt siRNA loci in *ddm1* leaf, as did shorter siRNAs (**Fig. 5C and Supplemental Fig. S14**). Taken together, these analyses reveal that the *ddm1* mutant causes leaf to resemble gametes in terms of siRNA expression, and suggest that loci that gain siRNA expression in either gamete are targets of methylation by DDM1, but not DRM2, in vegetative tissues.

## DISCUSSION

The male and female gametes of flowering plants are highly dimorphic, which is reflected in their very divergent transcriptomes (Anderson et al. 2013), as well their chromatin and histones (Ingouff et al. 2010), suggestive of major differences in reprogramming of their genomes prior to fertilization. The results from this study show that the small RNA transcriptomes of the rice gametes are very different from each other, as well as from vegetative cells. While both sperm and egg had reduced abundance of miRNAs relative to siRNAs, each gamete expressed a diverse set of miRNAs (**Supplemental Fig. S1**). Nevertheless, in both gametes, the most abundant miRNAs were from the miR159 family, with egg being more extreme (76% of total miRNAs; **Fig. 1**). miR159 targets a family of R2R3 myb domain transcription factors, and paternally expressed miR159 has been found to be important for regulation of endosperm development (Zhao et al. 2018), but as yet no biological functions have been reported for maternal miR159. The similarities between miRNA expression patterns between egg and ovary might be indicative of cell-to-cell miRNA mobility, but any such miRNA mobility is unlikely to be general, because ovary-specific miRNAs such as miR166a are not detectable or at very low levels in the egg cell (**Fig. 1C**). In Arabidopsis, expression of miR845 in pollen leads to production of *Gypsy* and *Copia* siRNAs called epigenetically activated siRNAs, or easiRNAs (Creasey et al. 2014; Borges et al. 2018; Martinez et al. 2018). However, we could not find any annotated miRNA or any genomic locus with significant homology to miR845 in rice, suggesting that *Gypsy* siRNAs in rice gametes use a different pathway. Expression of 24nt and 21nt phasiRNAs clearly differentiated sperm and egg, respectively (**Fig. 2**). 24nt phasiRNAs are expressed in meiotic anthers in diverse angiosperms including rice (Johnson et al. 2009; Zhai et al. 2015b; Xia et al. 2019). In female reproductive tissues, phasiRNAs are not well characterized, but an argonaute called MEL1 that binds 21nt phasiRNAs is required for both male and female meiosis (Nonomura et al. 2007; Komiya et al. 2014).

The most striking feature of both gametes was the depletion of 24nt siRNAs from heterochromatin boundaries (also called mCHH islands, **Fig. 6**) where they are found in typical vegetative tissues (Gent et al. 2013; Li et al. 2015), constituting key components of canonical RdDM [reviewed in (Cuerda-Gil and Slotkin 2016)]. In spite of this loss of 24nt siRNAs from canonical RdDM loci, both sperm and egg had similar abundance of 24nt siRNAs relative to other siRNA lengths but each had a distinct genomic distribution of 24nt siRNAs (**Fig. 2 and 3**). The redistribution of 24nt siRNAs into heterochromatin in sperm might be related to sperm-specific chromatin modifications, which could facilitate access of RNA pol IV to repetitive loci that would normally be inaccessible. Alternatively, the sperm siRNAs could be transferred from the pollen vegetative cell, which is known to lose heterochromatic modifications in Arabidopsis (Mérai et al. 2014). Dramatic changes in chromatin structure also occur in the production of female reproductive cells in plants, which may lead to corresponding changes in small RNA expression or activity [reviewed in (Wang and Köhler 2017)]. It is tempting to speculate that the loss of siRNAs from canonical RdDM loci in egg may be related to a loss of heterochromatin and hence heterochromatin boundaries in egg, but this explanation would also require either that the timing of loss of heterochromatin occurs earlier in the female germline during ovary development or occurs in parallel with egg in other ovary cells to explain the similarities between egg and ovary 24nt siRNAs. Alternatively, the chromatin status of egg may have more to with chromatin in ovary than in egg. While it has been proposed that siRNAs are translocated from the central cell into the egg cell (Ibarra et al. 2012), the similarities between ovary and egg raise the possibility that 24nt siRNAs in the egg cell might be translocated from the female reproductive tissues of the ovary.

**Figure 6.**
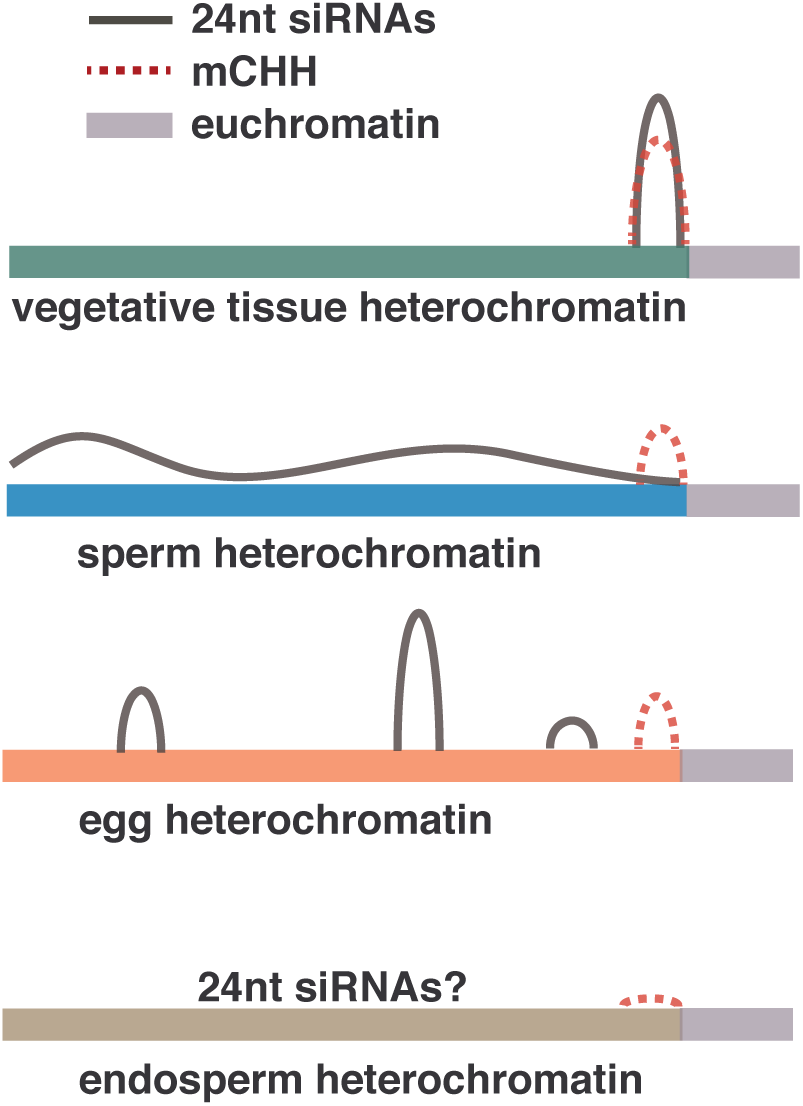
Schematic representation of 24nt siRNAs and CHH methylation in gametes.

A recent report on DNA methylation in rice pollen utilizing mutants in DNA glycosylase ROS1, revealed an intriguing correlation between demethylation in the pollen vegetative cell and increased methylation in sperm (Ibarra et al. 2012; Kim et al. 2019), which is consistent with previous findings in Arabidopsis (Ibarra et al. 2012). In this study, comparison of rice sperm cell PBAT methylomes with a previously published rice egg cell PBAT methylome (Park et al. 2016), and methylomes of vegetative tissues by conventional WGBS methods, detected different magnitudes of DNA methylation across these cell types, described as methylation reprogramming in the gametes (Park et al. 2016; Kim et al. 2019). We find from analysis of the published data from Kim et al. (2019) that they are consistent with our PBAT methylation libraries in this study, but that the deduced differences in methylation in sperm and egg are subtle in comparison to the differences in the 24nt siRNAs (**Fig. 3** *vs.* **Fig. 4**). More specifically, our analysis also found magnitudes of DNA methylation to be different between the different cell types, but the overall genome-wide patterns remained largely unchanged in all wild-type tissues we examined except mature endosperm (**Fig. 4**; summarized in **Fig. 6**). We propose that the gametes undergo a major re-programming of 24nt siRNAs, which is not correlated with a similar re-programming of DNA methylation. Since mCHH requires 24nt siRNAs, one explanation for the presence of mCHH at heterochromatin boundaries without 24nt siRNAs might be the removal of siRNAs after deposition of DNA methylation. It is also possible that mCHH was inherited from progenitor cells, in which case immediate progenitor cells would have had double the levels of mCHH. In this model, the generally low level of mCHH in gametes could be consistent with dilution of mCHH over one or more cell divisions.

DNA methylation in *drm2* mutants confirmed that mCHH and 24nt siRNAs in vegetative tissues defined canonical RdDM loci, as they were strongly dependent upon DRM2 for mCHH. The siRNA expression patterns of neither sperm nor egg could be explained by simple loss of 24nt siRNAs from canonical RdDM loci because 24nt siRNAs remained abundant relative to other siRNA lengths and even increased in abundance relative to miRNAs. In addition, removal of 24nt siRNAs from canonical RdDM loci in silico and analysis of the residual ones in seedling shoot yielded a genomic distribution unlike either gamete (**Supplemental Fig. S3B**). Thus, the pattern in both egg and sperm require both loss of 24nt siRNAs from canonical RdDM loci and gain of 24nt siRNAs at novel loci that do not undergo RdDM. In this respect, gametes share similarities with *ddm1* mutants in Arabidopsis, tomato, rice, and maize (Creasey et al. 2014; McCue et al. 2015; Corem et al. 2018; Long et al. 2018; Tan et al. 2018) and with chromomethylase mutants in maize (Fu et al. 2018). DDM1 facilitates methylation of heterochromatic DNA and has no known direct function in RdDM (Zemach et al. 2013; Lyons and Zilberman 2017). Our analysis of published *ddm1* mutant rice data (Tan et al. 2016; Tan et al. 2018) revealed that the loci with novel 24nt siRNA expression in either gamete tended to also gain siRNAs in *ddm1* mutant leaves, not just 24nt siRNAs but also 21 to 23nt siRNAs (**Fig. 5**). Both mCHH and 24nt siRNAs in *ddm1* mutant leaves were reduced at canonical RdDM loci, as previously reported. The effect *of ddm1* double mutant on siRNAs in leaves was not as striking as the difference between wild-type leaf and gametes, but the mutants clearly shifted leaf toward a more gamete-like profile. In maize *ddm1* mutant embryos, the effect is much stronger than in *ddm1* mutant rice leaf and more similar to rice gametes in that 24nt siRNAs are nearly completely lost from canonical RdDM loci (Fu et al. 2018).

The finding that a transgenic DDM1::GFP construct under the native DDM1 promoter expresses in Arabidopsis sperm but not in pollen vegetative cell supports a model in which absence of DDM1 in the pollen vegetative cell facilitates production of 21nt siRNAs that are transferred into sperm (Slotkin et al. 2009). While it is not clear how high levels of mCHG and mCG in pollen vegetative cell are produced in the absence of DDM1 (Calarco et al. 2012; Ibarra et al. 2012), it is clear that pollen vegetative cell undergoes major loss of heterochromatin in Arabidopsis (Mérai et al. 2014). The mRNA expression data in rice (Anderson et al. 2013) do not clearly indicate the DDM1 status in the pollen vegetative cell as it has reduced but not absent mRNA expression (**Supplemental Fig. S7**). It is likely though that the pollen vegetative cell in rice also loses heterochromatin, but with a stronger effect on 24nt siRNAs than on 21nt siRNAs. Regardless of the origin of 24nt siRNAs in rice gametes, their lack of correlation with mCHH in these cells raises two intriguing possibilities. The first possibility is that they direct chromatin modifications other than DNA methylation in gametes. The second is that the gametic siRNAs are primed for RdDM in the developing embryo. In this case, one or more RdDM factors missing in each gamete might become available upon fertilization and allow for immediate activity at loci not usually targeted in vegetative tissues.

## METHODS

### Rice stocks and growth conditions and gamete isolation methods

Rice (*Kitaake* variety) was grown in a greenhouse under nature light condition. Plants were irrigated with deionized water twice a week and supplemented with fertilized water every other week. Gametes were isolated as described (Anderson et al. 2013; Li et al. 2019). Briefly, ovaries were dissected from pre-anthesis flowers. A transverse cut was made at the middle region of the ovary. The lower part of the cut ovary was gently pushed by an acupuncture needle under a phase inverted microscope. Once the egg cell floated out from the ovary incision, it was captured by a fine capillary and frozen in liquid nitrogen. For small RNA, 35 – 50 cells were used as a biological replicate, and six biological replicates were collected. For PBAT libraries, 100 egg cells were used for each biological replicate. Sperm cells were collected by a Percoll gradient based method. Pollen grains were first released from mature flowers in a blender, then washed and osmotically burst. Pollen lysate was subject to a Percoll gradient centrifuge and sperm cells were collect between two density layers. Sperm-depleted pollen grains were taken as the pollen vegetative cell samples. For both small RNA and PBAT libraries, around 50 panicles with mature flowers were used for each biological replicate for sperm or pollen. Three replicates were collected for pollen vegetative cells for bisulfite sequencing. Eight biological replicates were collected for sperm cells for small RNA, and one more for bisulfite-seq. Ovaries were dissected from pre-anthesis flowers. For both small RNA and bisulfite sequencing, five ovaries were pooled to make a biological replicate. Three biological replicates were collected for ovaries for small RNA, and six more were collected for WGBS. Seedling shoot segments were collected from 7-day-old water-germinated rice seeds. For each seedling, the shoot was cut into small segments and immediately frozen in liquid nitrogen. One seedling was used for each biological replicate. Four biological replicates were collected for small RNA. Mature endosperm and embryo were separated after dry seeds were soaked in 6% NaOH in water at 57°C for 8 min, and pericarps were removed with forceps. The tissues were ground to a powder with a mini pestle in a 2ml Eppendorf tube, and DNA was extracted from each sample with a DNeasy Plant Mini Kit (Qiagen, 69104) and genotyped using the primers DRM2-Ri (TCTCACTACAAAGGCACCATAAAG) and DRM2-48F(CGAGGAGGAGGATGATACTAT) for the 48bp deletion allele and primers DRM2-Ri and DRM2-52F (CGAGGAGGAGGATGATATG) for the allele 52bp deletion allele.

### Generation of *drm2* mutation using CRISPR-Cas9 and genotyping

We used CRISPR-Cas9 to generate targeted mutations in the rice DRM2 gene (MSU: *LOC_Os03g02010)*. Single-guide RNA (sgRNA) sequences 5’-GGAGGAGGATGATACTAATT-3’ and 5’-GACAGGACTCCTCACTCTGA-3’, respectively, were designed by using the web tool https://www.genome.arizona.edu/crispr/ (Xie et al. 2014). CRISPR construct assembly was performed as described preciously (Khanday et al. 2019). Rice transformation was performed at UC Davis Plant Transformation Facility. One transgenic line carrying one in-frame 48bp deletion and one frame-shift 52bp deletion was selected for further experimentation. Since homozygous rice *drm2* mutants are sterile (Moritoh et al. 2012), we maintained the *drm2* mutation in a segregating population with the in-frame 48bp deletion. The 48bp allele was functional, as the plants carrying one or both 48bp alleles were phenotypically indistinguishable from wild-type. Genotyping was performed using two forward primers, F48: 5’-CGAGGAGGAGGATGATACTAT-3’ and F52: 5’-CGAGGAGGAGGATGATAT-3’, each specifically amplifies the −48bp or −52bp alleles, specifically, with one reverse primer: 5’-TCTCACTACAAAGGCACCATAAAG-3’. To genotype each sample, two separate PCR reactions were performed: F48 + R and F52 + R, at 59°C annealing temperature, 30 PCR cycles, and expected amplicon sizes were ∼500bp.

### RNA extraction and small RNA library construction

RNA extractions were performed using Ambion RNaqueous Total RNA kit (AM1931). We also perform an on-column DNase treatment using Qiagen DNase (79254). Total RNA was run on a Bioanalyzer to check for RNA integrity, under Eukaryotic total RNA-pico program. RNA input for egg cell was around 30 ng total RNA; 1 ng for sperm cells, 50 ng for ovaries, and 20 ng for seedlings. Small RNA libraries were made using NEXTflex Small RNA-seq kit v3 (NOVA-5132-05), with the following modifications. Since RNA input was low, a 1/4 dilution of adapters was used. The 3’ adapter ligation step was performed in 20°C for overnight. Sperm samples were amplified with 25 PCR cycles. All other samples were amplified with 15 – 20 cycles, except one of the four seedling replicates was amplified with 25 cycles. After amplification, libraries were run on a Bioanalyzer DNA High Sensitivity Assay. Libraries with 130 bp peak (adapter dimer peak) > 10% of the 150 pb peak (library product peak) were ran on a 10% TBE-Acrylamide gel (100 V for 1 hr). Gels were stained and the area around 150 bp was excised and purified according to recommendations of the NEXTflex Small RNA-seq kit.

### Mock egg isolations and qPCR quantification

Ovaries were dissected as during egg isolation as described previously (Li et al. 2019). About 1 µL of cell-free solution was collected into an Eppendorf tube and immediately frozen in liquid nitrogen. Thirty collections were combined as a single replicate, and two independent replicates were collected. RNA-extraction and library construction were performed as described above. A strong library band could be seen on a Bioanalyzer gel for a positive control ovary sample, but no band for negative water control or mock samples (**Supplemental Fig. S2B**), indicating that the lack of a band is not due to failed library preparation reactions, but lack of sufficient input RNA. To produce a DNA standard for qPCR absolute quantification, the following primers were used: P5_F: 5’-AATGATACGGCGACCACCGACATGACATTGACTATAAGGATGACG-3’ and P7_R: 5’-CAAGCAGAAGACGGCATACGAGATCGAGGCCGATGCTATACTTT-3’, using a plasmid template (Khanday et al. 2018). The target amplicon was 153bp and 49% CG content. The standard was PCR amplified, gel purified and quantified using a Nanodrop spectrophotometer. qPCR was performed using SYBR Green Master Mix (Biorad 1725270), with Illumina P5 and P7 universal primers (P5: 5’-AATGATACGGCGACCACCGA-3’ and P7-5’-CAAGCAGAAGACGGCATACGAGAT-3’), 60°C annealing temperature, 30 sec elongation time and 35 cycles. Serial dilutions of the standard (10^-1 to 10^-7, ten-fold dilution each step, two technical reps each) were used to fit the standard curve (**Supplemental Fig. S2C**). A 1/10 dilution of each library (three technical reps each) was used as templates for qPCR in the same run. Number of molecules in each library was calculated using the standard curve (**Supplemental Fig. S2D**).

### Genome annotations

The Os-Nipponbare-Reference-IRGSP-1.0 reference genome was used for all analyses (Kawahara et al. 2013). MSU7 Rice gene annotations were extracted from the all.gff genome annotation file downloaded from http://rice.plantbiology.msu.edu/pub/data/Eukaryotic_Projects/o_sativa/annotation_dbs/pseudomolecules/version_7.0/all.dir/. Genes that were flagged as transposons were removed, leaving a set of 39,953 genes. Transposons were annotated using RepeatMasker version 4.0.05 (http://www.repeatmasker.org/), parameters as follows: “-gff-species rice -s -pa 8”. miRNA annotations were downloaded from miRBase version 22 (Kozomara et al. 2019). To identify locations of the tandem repeat *CentO*, a consensus sequence (Zhang et al. 2013) was aligned to the genome using using Bowtie2 version 2.3.4.1 (Langmead and Salzberg 2012) parameters “-- local -a -f”. All alignments that were 100 or fewer basepairs apart were merged using BEDTools merge (Quinlan and Hall 2010). Locations of 5S rRNA and tRNA repeats were identified in the same way as *CentO*, but, using the GenBank reference sequence KM036285.1 for 5S rRNA and a set of tRNA sequences from The tRNAscan-SE Genomic tRNA Database for tRNAs (http://gtrnadb.ucsc.edu/GtRNAdb2/genomes/eukaryota/Osati/Osati-tRNAs.fa.) (Chan and Lowe 2016). For NOR annotation, an 18S ribosomal RNA gene, internal transcribed spacer 1, 5.8S ribosomal RNA gene, internal transcribed spacer 2, and 26S ribosomal RNA gene complete sequence from GenBank (KM036285.1) was aligned to the genome with Bowtie2 version 2.3.4.1, parameters “--local --ma 1 --mp 24,8 --rdg 20,48 --rfg 20,48 -a f”. All alignments that were 100 or fewer basepairs apart were merged using BEDTools.

### Small RNA sequencing analysis

Small RNA-seq reads were quality filtered and trimmed of adapters and of the four random nucleotides on the ends using cutadapt (Martin 2011) parameters “-u 4 -q 20 -a TGGAATTCTCGGGTGCCAAGG -e .05 -O 5 --discard-untrimmed -m 24 -M 29”, followed by a subsequent run with just “-u -4”. For previously published *ddm1, dmr2*, and root small RNA reads, which did not have four random nucleotides on the ends (Tan et al. 2016; Shin et al. 2018; Tan et al. 2018), parameters were “-q 20 -a TGGAATTCTCGGGTGCCAAGG -e .05 -O 20 -m 20 -M 25”. Reads were aligned to the genome with BWA-backtrack (version 0.7.15) (Li and Durbin 2009), parameters “aln -t 8 -l 10.” A single mapped position was kept per input read, regardless of the possibility of mapping to multiple locations. Except where indicated otherwise, multi-mapping reads were included in all analyses. Locations of 21nt and 24nt phasiRNA loci were identified using PHASIS version 3.3 (https://www.biorxiv.org/content/10.1101/158832v1). The phasdetect module was run on the set of small RNA read samples, followed by the phasemerge module, parameters “-mode merge -pval 1e-5” using each length from 20 to 25nt, but only 21 and 24nt lengths produced detectable phasiRNA loci (12 21nt phasiRNA loci and 67 24nt phasiRNA loci, **Supplemental Table S4**). The complete set of read alignments was compared with miRNA, phasiRNA, tRNA, 5S rRNA, and NOR RNA loci in the genome and all reads that aligned by at least 90% with any of these was categorized as such using BEDTools intersect. All other reads were categorized as siRNA reads and used for subsequent siRNA analyses. The uniquely mapping subset of siRNAs was defined by having MAPQ values of at least 20 using SAMtools [Li et al (2009)]. For all other analysis of overlaps with siRNAs and genetic elements (*Gypsy* retrotransposons, the *CentO* centromeric tandem repeat, Terminal Inverted Repeat (TIR) DNA transposons, and 24nt siRNA loci) we only counted siRNAs that overlapped by at least 50% of their lengths. CACTA elements were excluded from the TIR DNA transposons (**Supplemental Table S5**). Whole-genome small RNA heatmaps were made on 50Kb intervals using IGVtools (Thorvaldsdóttir et al. 2013). For better visualization of midrange values, heatmap intensity was maxed out at 7X coverage (per 50 million 24nt siRNAs). To identify 24nt siRNA loci, reads alignments were subsampled then combined from each sperm sample, from each egg samples, and from each seedling shoot sample to get as equal a representation as possible from each sample and a final combined number of 50 million in each using SAMtools view –s followed by SAMtools merge. Read coverage over 100bp non-overlapping loci was measured using BEDTools coverage. All loci that were overlapped by at least 25 24nt siRNAs that spanned at least 50 bp of the 100 bp were categorized as 24nt siRNA loci. 5’ nucleotide frequencies were calculated with FastQC, version 0.11.8. (https://www.bioinformatics.babraham.ac.uk/projects/fastqc/).

### miRNA expression analyses

miRNA expression data were organized into a matrix, with each row as an individual miRNA and each column as a library (**Supplemental Dataset 1**). R package Edge R was used to analyze miRNA expression (McCarthy et al. 2012). miRNAs were normalized by total small RNA, and filtered for >1 counts per million reads (CPM). Libraries were then further normalized by the TMM method (Robinson and Oshlack 2010), as recommended by the EdgeR package. Differential expression analyses were performed under log2FC > 1 and FRD < 0.05 cutoffs. Differential expressing miRNA were visualized under counts per million miRNAs. Principal component analyses were performed using log-transformed CPM values. Clustering analyses (**Supplemental Table S3**) were performed using hierarchical clustering, and assignment of miRNA into clusters were done using the Dynamic Tree Cut R pacakge (Langfelder et al.2008).

### Preparation of WGBS libraries

PBAT libraries were prepared using Pico Methyl-Seq Library Prep Kits (Zymo D5456). Sperm and egg isolates of approximately 100 cells each were diluted in 200 ul of 10 mM Tris, pH 8 in a 1.5 ml tube, then centrifuged for 10 minutes at 16,000 x G at 4°C. The supernatant was removed except for 9 ul at the bottom of the tube. The 9 ul was pipetted up and down 10 times, then transferred to a 0.2 mL tube. 10 ul M-Digestion Buffer and 1ml Proteinase K (Zymo D3001-2-5) were added and incubated for 20 minutes at 50°C. For pollen vegetative cells, 12 ul was retained after the initial dilution in Tris buffer, and 13 ul Zymo M-Digestion Buffer and 1ml Zymo Proteinase K (20 mg/mL) were added. Following incubation for 20 minutes at 50°C, the pollen vegetative cells were centrifuged again for 5 minutes at 16,873 x G, then 20 ul of supernatant transferred to a new tube. Subsequent steps were as directed in the Pico Methyl-Seq Library Prep Kit protocol, version 1.2.0, with a 16-minute incubation in L-Desulphonation buffer, and 5:1 ratios of DNA binding buffer in DNA purification steps. Conventional WGBS libraries from endosperm, embryo, and ovary, were prepared using the methylC-seq method (Urich et al. 2015).

### WGBS analysis

MethylC-seq and PBAT reads were quality filtered and trimmed of adapters using cutadapt [Martin (2011)], parameters as follows: “-q 20 -a AGATCGGAAGAGC -e .1 -O 1 -m 50”. PBAT reads were aligned to the genome with BS-Seeker2 (version 2.1.5) (Guo et al. 2013) with parameters as follows: “--aligner=bowtie2 --bt2--end-to-end -m 1 -t Y -s 5 -e 100” methylC-seq reads were aligned similarly, but BS-Seeker2 parameters were modified to “-m 1 -- aligner=bowtie2”. For all reads except paired-end, previously published reads (Tan et al. 2016), PCR duplicates were removed prior to alignment with the BS-Seeker2 FilterReads.py module. DMRs were identified with CGmapTools version 0.1.1 (Guo et al. 2018). Biological replicates were first merged using the CGmapTools mergelist tosingle module, then the set of cytosines with coverage in both samples were identified using the CGmapTools intersect module. Methylation comparisons were made using the CGmapTools dmr module, parameters “-c 3 -C 50 -s 100 -S 100 -n 5”. DMRs were selected based on four criteria: at least five measured cytosines in the region; a P-value less than 0.001; absolute difference in methylation proportion of greater than 0.25 (wild-type value minus *drm2* mutant value), and a relative difference in methylation proportion of less than 0.3 (wild-type value divided by *drm2* mutant value).

### mRNA expression analysis

Previously published mRNA reads [Anderson et al (2013)] were aligned to the genome using Tophat, version 2.0.13 [Kim et al (2013)], parameters as follows: “--read-realign-edit-dist=0 --min-intron-length=15 --max-intron-length=20000 --max-multihits=1 --microexon-search --library-type=fr-unstranded --b2-very-sensitive”. For uniquely mapping reads only, “-- prefilter-multihits” was also included. The number of reads that overlapped with genomic features was counted using BEDTools intersect, requiring that half of each read’s mapped length overlapped with a feature using the “-f .5” parameter to be counted.

All R-script for statistical analyses and data visualization can be found at: https://github.com/cxli233/gamete-small-RNA

## DATA ACCESS

All small RNA and WGBS data have been deposited in the Sequence Read Archive, BioProject PRJNA533115.

## Supplemental Figures and Tables

**Supplemental Fig. S1:**

Differential miRNA expression and clustering

**Supplemental Fig. S2:**

Mock egg isolations and qPCR quantification of mock small RNA libraries

**Supplemental Fig. S3:**

Genome-wide view of 21nt, 22nt, and 24nt siRNAs

**Supplemental Fig. S4:**

Proportion of siRNAs per each specific length at sample-specific or intersection loci

**Supplemental Fig. S5:**

Transposon content of sample-specific 24nt siRNA loci and distances to nearest genes

**Supplemental Fig. S6**:

Highly-expressed sperm genes (>10 TPM) are also depleted of flanking 24nt siRNAs

**Supplemental Fig. S7**:

Expression of RdDM and methylation related factors in gametes

**Supplemental Fig. S8**:

Proportion of mRNA transcripts mapping to transposons in gametes

**Supplemental Fig. S9**:

Scatter plot of individual transposon copy transcripts vs. 24nt siRNAs

**Supplemental Fig. S10:**

Genome-wide view of DNA methylation

**Supplemental Fig. S11:**

Methylation metaplots of complete set of PBAT libraries analyzed

**Supplemental Fig. S12:**

DNA methylation of 24nt siRNA loci

**Supplemental Fig. S13:**

Metaplot for 24nt siRNAs in *ddm1* and *drm2* leaf from Tan et al. (2016)

**Supplemental Fig. S14:**

Proportion of siRNAs per each specific length at sample-specific and intersection loci from leaf (Tan et al 2016).

**Supplemental Table S1**:

List of all small RNA libraries, read counts, and SRA accession numbers

**Supplemental Table S2**:

List of all WGBS libraries, read counts, and SRA accession numbers

**Supplemental Table S3**:

Assignment of miRNAs into clusters

**Supplemental Table S4:**

List of phasiRNA loci

**Supplemental Table S5:**

Small RNA composition organized in a spreadsheet

**Supplemental Dataset 1:**

Table of miRNA raw counts for all small RNA libraries

## Acknowledgements

We would like to thank Daniel Jones for assistance in sperm isolation; Zach Liechty, Christian Santos and Joseph Edwards for assistance in R programming; and Alina Yalda and Jake Anichowski for greenhouse maintenance. The UC Davis Genome Center provided Illimuna sequencing services. This research was funded by the National Science Foundation (IOS-1547760) and the USDA Agricultural Experiment Station (CA-D-XXX-6973-H).

